# Ciliopathy interacts with neonatal anesthesia to cause non-apoptotic caspase-mediated motor deficits

**DOI:** 10.1101/2024.11.27.624302

**Authors:** Nemanja Sarić, Zeynep Atak, Courtni Foster Sade, Nikita Reddy, Gabrielle Bell, Christina Tolete, May T. Rajtboriraks, Kazue Hashimoto-Torii, Vesna Jevtović-Todorović, Tarik F. Haydar, Nobuyuki Ishibashi

## Abstract

Increasing evidence suggests that anesthesia may induce developmental neurotoxicity, yet the influence of genetic predispositions associated with congenital anomalies on this toxicity remains largely unknown. Children with congenital heart disease often exhibit mutations in cilia-related genes and ciliary dysfunction, requiring sedation for their catheter or surgical interventions during the neonatal period. Here we demonstrate that briefly exposing ciliopathic neonatal mice to ketamine causes motor skill impairments, which are associated with a baseline deficit in neocortical layer V neuron apical spine density and their altered dynamics during motor learning.. These neuromorphological changes were linked to augmented non-apoptotic neuronal caspase activation. Neonatal caspase suppression rescued the spine density and motor deficits, confirming the requirement for sublethal caspase signaling in appropriate spine formation and motor learning. Our findings suggest that ciliopathy interacts with ketamine to induce motor impairments, which is reversible through caspase inhibition. Furthermore, they underscore the potential for ketamine-induced sublethal caspase responses in shaping neurodevelopmental outcomes.

## Introduction

Congenital heart disease (CHD) is the most common birth defect and is associated with an increased risk for a wide range of neurodevelopmental and behavioral problems. CHD patients frequently exhibit motor skill deficits with estimates of more than half of them having moderate to severe motor development issues(*1, 2*). The etiology of these neurological impairments is cumulative and multifactorial, including intrinsic genetic predisposition and stressors such as neonatal cardiac surgery.

Forward recessive genetic screening performed in mouse fetuses revealed a compelling association between genes encoding proteins involved in ciliogenesis, ciliary signaling pathways and trafficking and CHD incidence(*3*). These findings are corroborated by clinical data, where CHD patients show a high prevalence of *de novo* mutations in cilia-related genes and general ciliary dysfunction(*4, 5*). The primary cilium acts as a sensory organelle that regulates cellular signal transduction, thus modulating cell behavior and physiology in various tissues including brain and heart. Importantly, recent evidence implicates proteins involved in ciliary intraflagellar transport (IFT), the trafficking of proteins along the microtubular scaffold of the primary cilium, in the regulation of cellular apoptosis and mitochondrial stress responses(*6, 7*).

Anesthesia remains one of the most consequential and beneficial developments in modern medicine. Sedation with anesthetic agents such as propofol and ketamine is often employed in the care of children prior to catheter or surgical interventions, as well as some diagnostic procedures, necessary in neonates with structural defects such as CHD patients (*8, 9*). Despite these desirable properties, the last several decades of both preclinical and clinical research have demonstrated correlations and causal relationships between prolonged and/or repeated anesthesic and sedative exposure and developmental neurotoxicity(*10, 11*). Sublethal cellular responses to environmental stressors, which do not result in neuronal death but nevertheless alter neuronal cytoarchitecture and physiology, are only beginning to be investigated(*12*).

Many of the compounds used for sedation in major pediatric interventions including ketamine are pleiotropic in action, causing dose-dependent hypnotic, analgesic, and amnesic effects, mostly *via* their antagonism of N-methyl-D-aspartate (NMDA) and α-amino-3-hydroxy- 5-methyl-4-isoxazolepropionic acid (AMPA) receptor signaling in the brain(*13*). In addition to their dampening effects on excitatory neurotransmission, these compounds also have agonistic properties on γ-aminobutyric acid type A (GABAA) receptors(*14–16*). Mounting preclinical data from rodent and non-human primate studies have suggested that anesthesia exposure enhances neurodegeneration in the developing perinatal brain(*17, 18*). Indeed, clinical data found specific associations between increasing ketamine doses and greater motor impairments as assessed by Bayley III motor scores in infants with CHD at 18 months of age(*19*). The described convergence on impairments of the motor system between CHD and perioperative factors might result from independent effects of either factor or potentially, a synergistic interaction. Therefore, normal ciliary integrity and function might play a role in modulating neuronal apoptotic stress responses to sedative or anesthetic exposures, thus impacting neurodevelopmental outcomes in CHD patients undergoing surgical procedures during the early postnatal period.

*Ift88* is one of the candidate genes responsible for left ventricular outflow tract obstruction, conotruncal defects, and heterotaxy(*20*). Loss of function of the *Ift88* gene results in the failure of primary cilium synthesis and has previously been used to model primary ciliopathy in mouse thyrocytes, cranial neural crest, endothelial cells, and the inner ear(*21–24*). Targeted *Ift88* loss of function in the developing murine heart demonstrated multiple CHD-associated heart defects, including double outlet right ventricle and atrioventricular septal defects(*25*). Prior work with a forebrain-specific *Ift88* mutant, driven by an *Emx1*-dependent Cre recombinase(*26*), found that perinatal ethanol exposure leads to acutely elevated caspase 3 activation in pyramidal neurons of the primary motor cortex associated with long-term degenerative changes in dendritic arborization(*27*). Given that ethanol acts as a potent antagonist of NMDA-mediated neurotransmission(*28*), it is plausible that general anesthetic and sedative agents such as ketamine might interact with the loss of cilia in a similar manner, augmenting the risk of motor impairments in patients with ciliary dysfunction such as those with CHD. To model the interaction between a sedative procedure and ciliary dysfunction we exposed the forebrain-specific *Ift88* mutant mice to ketamine, followed by characterization of their motor skill learning and neuronal morphology.

In this study we find that acute exposure to a single dose of ketamine during early postnatal development leads to enhanced caspase 3 activation in layer V motor cortical pyramidal neurons of a forebrain-specific model of ciliopathy (*Ift88* cKO). Neonatal ketamine exposure led to pronounced long-term fine motor skill impairments specifically in this group, which were associated with a reduction in dendritic spine density of the apical arbor of layer V motor cortical pyramidal neurons. In addition, the ketamine-treated *Ift88* cKO pyramidal neurons exhibited reduced dendritic arborization complexity in adulthood. Notably, treatment with a pan-caspase inhibitor QVD-Oph rescued the motor learning as well as spine deficits, consistent with a key role for caspase signaling in the neuronal and behavioral alterations observed in *Ift88* cKO animals. Caspase 3 activation after ketamine treatment did not lead to overt neuronal loss, arguing for a sublethal mode of augmented caspase function in this model.

## Results

### Caspase 3 activation in layer V ciliopathic motor neurons exposed to ketamine does not cause apoptosis

We initially set out to assess the potential for interaction between ketamine exposure and neuronal ciliary deficits during the early murine postnatal window. To determine the effect of neonatal ketamine on caspase activation in neurons of the ciliopathic neocortex, a single dose of ketamine (20 mg/kg) or vehicle was administered intraperitoneally to P7 pups (equivalent to human neonate) born to crosses between *Emx1-Cre; Ift88^fl/+^* and *Ift88^fl/f^* mice (Fig. 1A). This ketamine dose was chosen since prior work demonstrated it to be the minimal subanesthetic dose necessary for eliciting a subtle yet significant elevation in cleaved caspase 3 (CC3) immunoreactivity in the caudate putamen brain area, and deemed to be equivalent to a sedating/subanesthetic ketamine dose for an infant human(*29*). In this way we endeavored to test whether ciliopathy augments ketamine-induced CC3 activation. CC3 signal intensity was assessed in neocortical neurons at 16 hours following treatment, using signal co-localization between NeuN and CC3. We observed strongly elevated neuronal CC3 immunoreactivity in the ketamine-treated *Ift88* cKO group, forming a specific pattern across the perinatal neocortex (Fig. 1B-C). This result corroborated previous findings using ethanol-exposed *Ift88* mutant animals(*27*), which showed increased caspase 3 activation in neurons of the primary motor cortex (PMC), specifically in ciliopathic *Ift88* cKO neurons exposed to ethanol. Caspase 3 activation was enhanced in sparse populations of neurons in the medial prefrontal (mPFC) and primary somatosensory cortex (PSC), while strikingly elevated in deeper layers of the PMC and across the gustatory cortex (GC) (Fig. 1B). The pattern of elevated caspase activation in the ketamine treated ciliopathic group (*Ift88* cKO + KET) was not uniform among deep layer neurons with neighboring cells displaying contrasting levels of signal (Fig. 1C). This heterogeneity in caspase activation is intriguing as it suggests variable sensitivity to ketamine among neurons occupying the same cortical layers and/or functional specialization, regardless of their ciliary integrity.

**Figure 1.**
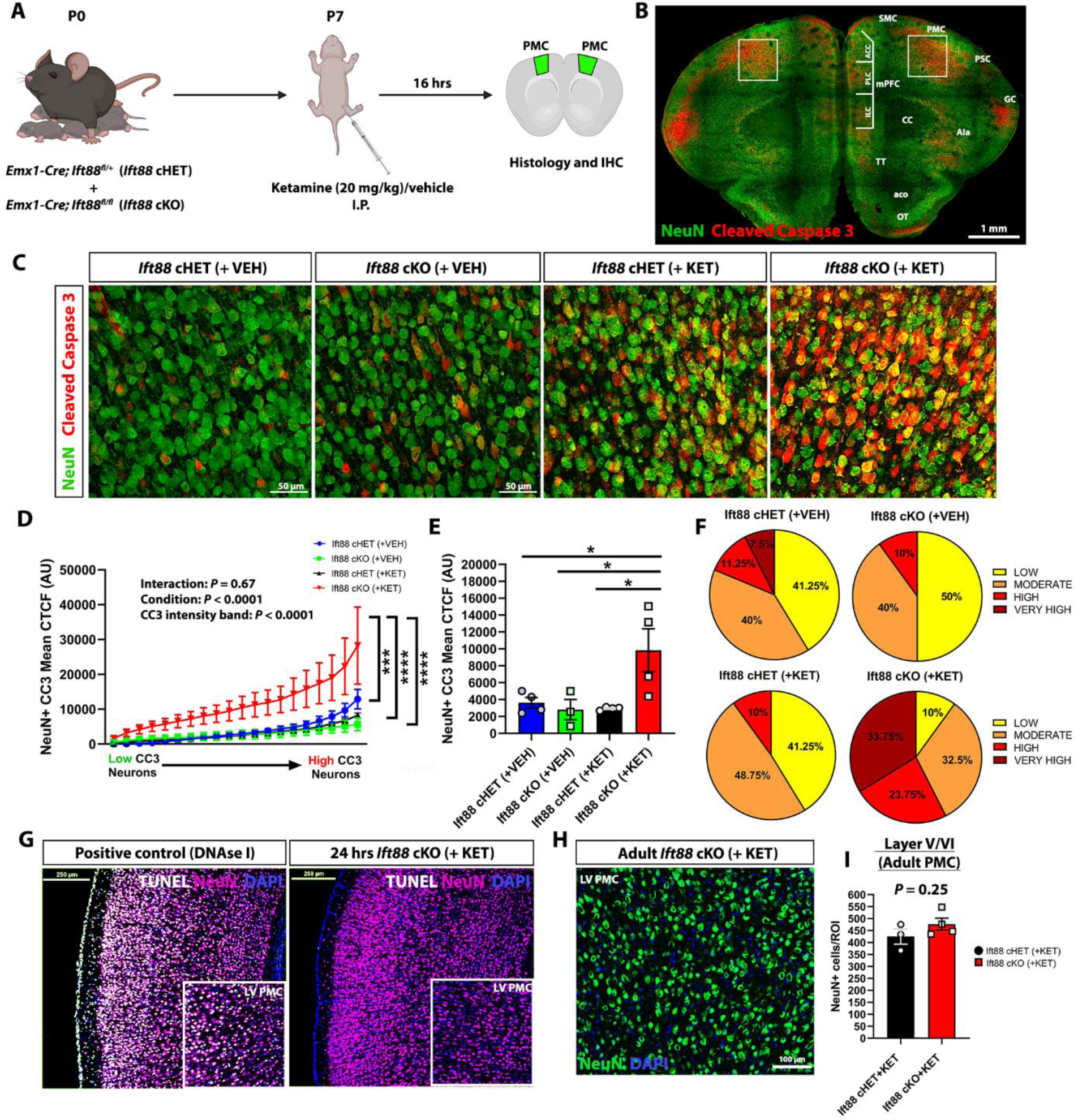
Enhanced non-apoptotic caspase 3 activation in ciliopathic motor cortex deep layer neurons following ketamine exposure. (**A**) Schematic illustrating ketamine treatment regime followed by immunohistochemical caspase signaling assessment at 16 hours post-exposure in *Ift88* mutant mice. (**B**) Representative cleaved caspase 3 (red) staining pattern in a coronal section of the anterior frontal cortex of ketamine treated *Ift88* cKO, with strong signal (white boxes) observed in deep layers (layer V/VI) of the primary motor cortex (PMC) among other regions. (**C**) Magnified micrographs show co-localization of cleaved caspase 3 signal with neuronal NeuN in layers V/VI of the PMC, and comparison between vehicle and ketamine treated *Ift88* cHET and cKO mice. (**D**) Line graph displaying mean cleaved caspase 3 (CC3) corrected total cell fluorescence (CTCF) values across 20 representative neurons/PMC region of interest (ROI) (ordered from neurons with lowest to neurons with highest mean CC3 CTCF), and their comparison between vehicle and ketamine treated *Ift88* cHET and cKO mice. Note enhanced CC3 signal specifically in the ketamine treated *Ift88* cKO group. (**E**) Quantification of the mean CC3 CTCF across sampled deep layer neurons from groups shown in **C-D**. (**F**) Pie charts showing proportions of neurons displaying low (0-2000), moderate (2000-6000), high (6000-10,000) and very high (10,000+) CC3 CTCF values across treatment groups. (**G**) Representative positive control (DNAse I treated) and *Ift88* cKO (ketamine treated) TUNEL staining at 24 hours post-exposure to ketamine. Note lack of detectable TUNEL signal in ciliopathic samples. (**H**) Representative micrograph of NeuN immunoreactive neurons from adult (8-10 weeks) *Ift88* cKO PMC layers V/VI. (**I**) Quantification of NeuN^+^ neurons in control and ciliopathic PMC deep layers. Note no difference in neuronal numbers. Values in **D-E** and **I** are mean ± standard error (SEM) (n=3-4 in each; 2 male + 1 female samples in Ift88 cKO (+VEH) group, 2 male + 2 female samples in other groups). \**P* < 0.05; \*\*\**P* < 0.001; \*\*\*\**p* < 0.0001. Data in (**D**) was analyzed using a 2-way ANOVA with post-hoc Tukey’s multiple comparisons test. Data in (**E**) was analyzed using a 1-way ANOVA with Tukey’s multiple comparisons test. Data in (**I**) was analyzed by an unpaired student’s t test. Scale bars: (**B**) – 1 mm; (**C**) – 50 µm; (**G**) – 250 µm; (**H**) – 100 µm. Abbreviations: I.P.-intraperitoneal, CC3-cleaved caspase 3, mPFC-medial prefrontal cortex, ILC-infralimbic cortex, PLC-prelimbic cortex, ACC- anterior cingulate cortex, SMC-secondary motor cortex, PMC-primary motor cortex, PSC-primary somatosensory cortex, GC-gustatory cortex, AIa-agranular insular cortex, CC-corpus callosum, OT-olfactory tubercle, aco-anterior commissure, TT-taenia tecta.

Quantification of mean CC3 corrected total cell fluorescence (CTCF) revealed a highly significant increase in caspase 3 activation among ciliopathic (*Ift88* cKO) ketamine-exposed deep layer neurons (Fig. 1D). Interestingly, while a higher CC3 signal was present in *Ift88* cKO neurons across a range of CC3 intensities, the neurons with higher CC3 CTCF values showed a highly statistically significant CC3 signal enhancement compared to neurons in other groups. This difference was also reflected in the mean CC3 CTCF value comparisons (Fig. 1E) as well as the proportions of low, moderate, or high CC3 intensity neurons present in layer V/VI primary motor cortex (Fig. 1F). Vehicle treated *Ift88* cHET and cKO mice did not show differences in CC3 intensity suggesting that the loss of primary cilia alone does not influence signaling through caspase 3 in perinatal neocortical neurons. On the other hand, ketamine treatment did not result in CC3 elevation in *Ift88* cHET neurons, indicating that a single dose of ketamine does not have an observable impact on caspase signaling in neurons expressing functional primary cilia.

Caspase 3 activation has been classically associated with the initiation of apoptosis and cell death, which culminates with the process of DNA fragmentation. To assess DNA fragmentation in ketamine treated *Ift88* cKO neurons we performed terminal deoxynucleotidyl transferase dUTP nick end labeling (TUNEL) staining at 24 hours following ketamine administration. We chose this time window based on prior research in rat models of focal cerebral ischemia, which showed substantial DNA fragmentation in the cortex and striatum at 24 hours post-reperfusion injury(*30*). Ciliopathic ketamine-exposed neocortical neurons did not display detectable TUNEL immunoreactivity, in stark contrast to DNAse I-treated positive control tissue (Fig. 1G). To follow up and confirm this finding, we examined neuronal numbers in PMC deep layers of adult (8-10 weeks) *Ift88* cHET and cKO ketamine treated animals (Fig. 1H). Quantification of NeuN immunoreactive cells in the deep layers of the PMC did not show a statistically significant difference in neuronal number between control and ciliopathic brains (Fig. 1I). A lack of neuronal loss in the ciliopathic PMC suggests that the augmented neuronal caspase signaling, observed shortly after ketamine exposure, does not lead to apoptosis and long-term cell loss. This finding agrees with a multitude of studies demonstrating non-apoptotic, sub-lethal roles for caspase signaling in both physiological and pathological contexts(*12*).

These results point to a potential causative correlation between loss of primary cilia and acute ketamine exposure in driving enhanced caspase 3 activation in neocortical deep layer motor neurons, whereas neither factor is sufficient on its own for the observed changes.

### Ciliopathic layer V motor neurons exposed to ketamine have reduced arbor complexity

Cleaved caspase 3 is known to destabilize microtubule networks in neurons, leading to reduced arborization and enhanced dendritic pruning(*31, 32*). To investigate whether the augmented caspase signaling observed following ketamine exposure influences dendritic arborization in our experimental groups, we crossed *Thy1-GFP* reporter mice with the *Emx1-Cre; Ift88^flox^*lines, followed by ketamine treatment at P7. Brain tissue was collected and histologically processed from animals that had undergone rotarod motor training and multiphoton imaging (Fig. 2A). Arborization complexity analysis using the Sholl method revealed a statistically significant reduction in branching complexity in ketamine exposed *Ift88* cKO pyramidal neurons (Fig. 2B- C). Analyses of neuronal soma size did not show clear differences by genotype, suggesting that only the ciliopathic dendritic compartment was affected by the ketamine treatment (Supplementary Fig. S1A-B). These data argue that caspase 3 activation reduces the branching complexity of ciliopathic layer V pyramidal neurons (Fig. 2C), signifying structural neuronal changes rather than apoptotic cell loss.

**Figure 2.**
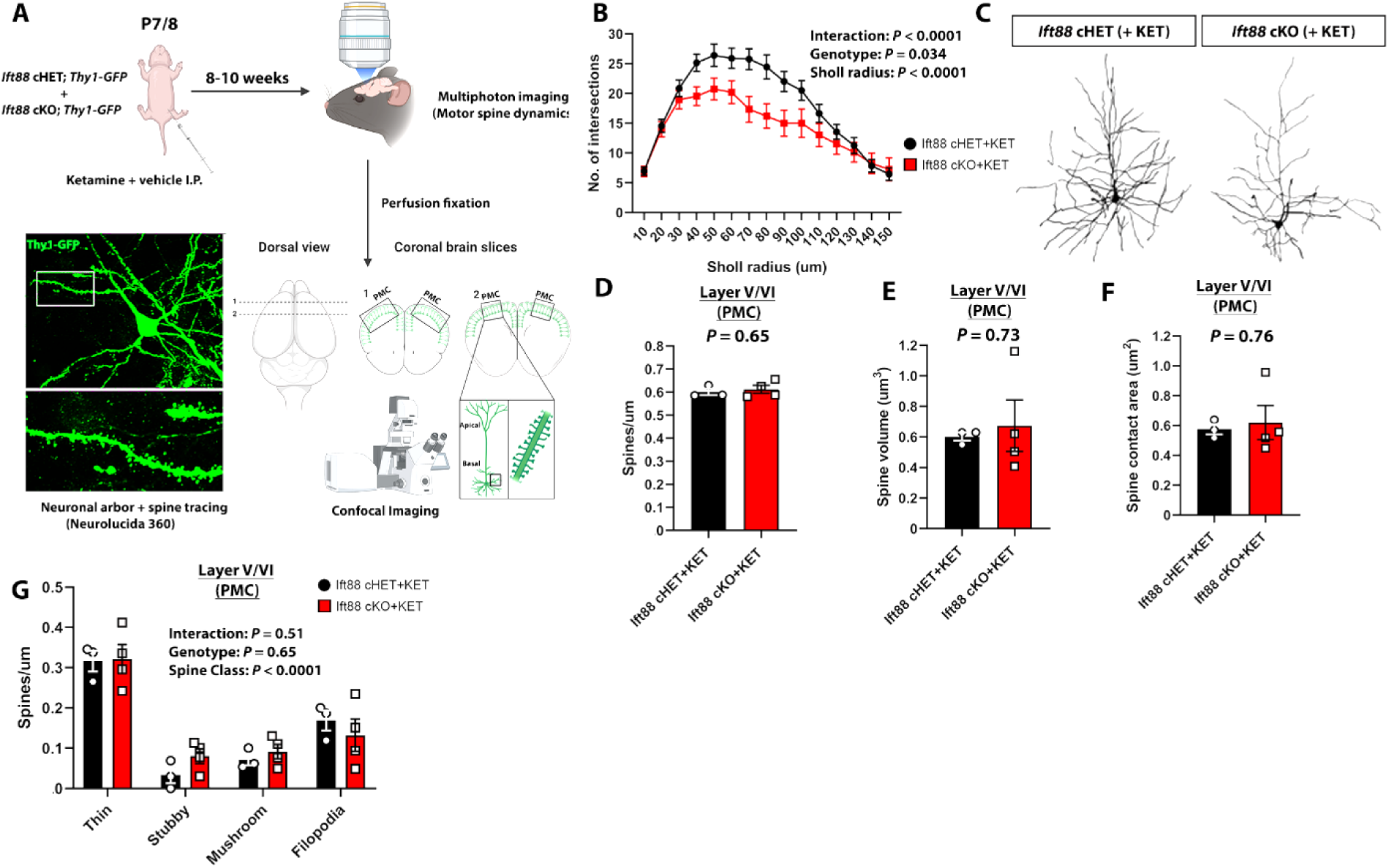
Ketamine treatment diminishes layer V neuronal arborization complexity in ciliopathic mice. (**A**) Schematic showing tissue processing regime for examining layer V PMC pyramidal neuron arborization and basal spine morphometry following multiphoton imaging. (**B**) Quantification of dendritic intersections, a proxy for arborization complexity, of layer V pyramidal neurons from ketamine treated controls (*Ift88* cHET) and ciliopathic (*Ift88* cKO) mice using Sholl analysis of dendrite traces obtained with Neurolucida 360. Note decreased complexity of ketamine treated *Ift88* cKO neurons. (**C**) Representative monochrome traces of *Thy1-GFP* reporter labeled layer V pyramidal neurons from control (*Ift88* cHET) and ciliopathic (*Ift88* cKO) mice. (**D-F**) Quantification of basal dendritic spine density (**D**), volume (**E**) and contact area (**F**). No differences were seen in parameters between groups. (**G**) Bar graphs showing breakdown of spine class densities in the basal arbors of layer V pyramidal neurons from the ketamine treated groups. Values in (**B**) and (**D**-**G**) represent mean ± standard error (SEM) (n=3 in **B**, 2 male + 1 female samples in each group; n=3-4 for **D**-**G**, 2 male + 1 female samples in *Ift88* cHET (+KET) group, 2 male + 2 female samples in *Ift88* cKO (+KET) group). \**P* < 0.05. The statistical test used for (**B**) was a 2- way Mixed Effects ANOVA, implemented as previously described(*65*). Data in (**D**-**G**) was analyzed by unpaired student’s t tests. Abbreviations: PMC-primary motor cortex, KET-ketamine, I.P.-intraperitoneal.

Caspase signaling within neurons is often accompanied by neuroinflammatory changes in the local microenvironment. One of the principal drivers of neuroinflammation are microglial cells(*33*), which infiltrate the injury site/s for the phagocytosis of dying cells and cellular debris and neuroprotection. Microglia have also been demonstrated to engage in presynaptic trogocytosis, a process of partial engulfment of intracellular material through cell-to-cell contacts(*34*). Despite a lack of conclusive evidence for a microglial role in alterations to postsynaptic spine elements, these previous findings argue that microglia do remodel synaptic contacts in the postnatal brain. Thus, we examined microglial numbers in the deep layers of the PMC corresponding to the areas where CC3^+^ neurons were detected, using Iba1 immunoreactivity in the ketamine treated *Ift88* cHET and cKO groups (Supplementary Fig. S2A). We did not detect statistically significant differences in the total numbers of microglia between groups, suggesting that neonatal ketamine treatment does not elicit increased microglial infiltration in the ciliopathic group (Supplementary Fig. S2B).

Caspase signaling can elicit changes in neuronal dendritic spines alongside alterations to the arborization pattern. To establish whether perinatal ketamine affects the basal dendritic compartment, we performed spine tracing, morphometry and classification on basal dendritic branches imaged from the same samples obtained for dendritic arborization studies (Fig. 2A). We did not detect differences in mean basal spine density (Fig. 2D), spine volume (Fig. 2E) or spine contact area (Fig. 2F) between control and ciliopathic ketamine-treated groups. In addition, basal spine classification and comparison did not show significant changes in the density of individual immature and mature spine classes (Fig. 2G), although there was a trend towards an increase in stubby spine density in the *Ift88* cKO group.

In summary, our morphometric analyses of layer V pyramidal neurons revealed a reduction in arborization complexity of ciliopathic neurons following ketamine exposure, without observable differences in neuronal cell numbers or basal spine density and morphology. This finding suggests that ketamine-induced caspase signaling results in long-term remodeling of basal dendrites of maturing layer V ciliopathic pyramidal neurons.

### *Ift88* cKO mice exposed to ketamine display fine motor deficits

Extensive non-apoptotic caspase 3 activation was observed in cilia-deficient motor cortical neurons and was associated with morphological changes. To investigate the functional consequences of the observed increase in neuronal caspase signaling after ketamine treatment in *Ift88* cKO animals, we assessed fine motor skill acquisition in our experimental cohorts. Fine motor skill development is known to be deficient in individuals with CHDs after neonatal cardiac surgery(*35, 36*). Forelimb extension and food pellet reaching is a robust and well-validated fine motor skill assay in rodents(*37*). *Ift88* cHET and cKO mice treated with either vehicle or ketamine at P7 were left to mature prior to fasting at 8-10 weeks of age to ensure adequate food motivation (Fig. 3A).

**Figure 3.**
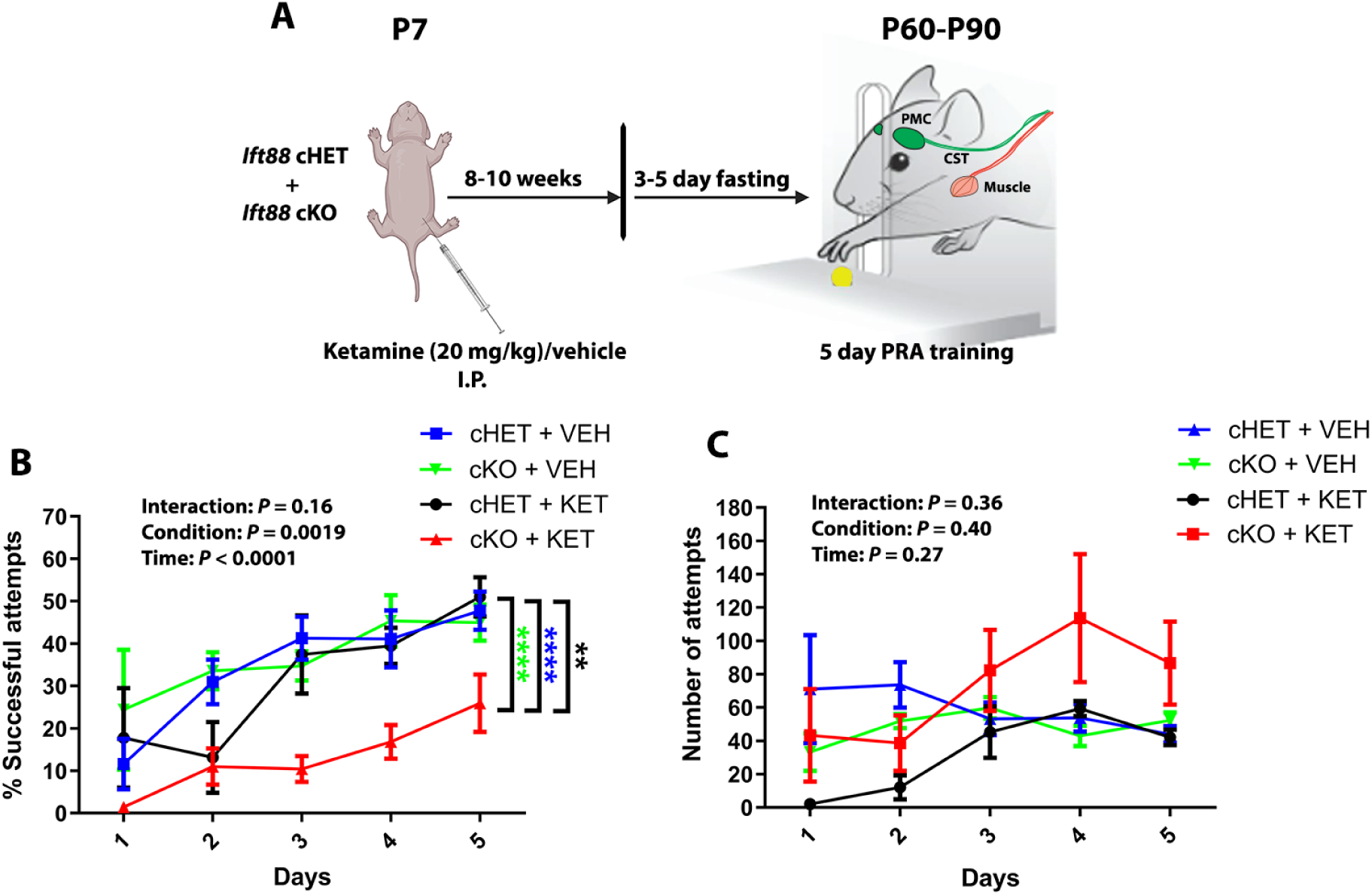
Ketamine exposed *Ift88* cKO mice display fine motor skill deficits. (**A**) Diagram representing treatment regime and timeline for assessment of pellet reach fine motor skills in control (*Ift88* cHET) and mice with forebrain-specific ciliopathy (*Ift88* cKO). Numbers used: vehicle treated *Ift88* cHET, *n* = 5 (3 males + 2 females); vehicle treated *Ift88* cKO, *n* = 4 (2 males + 2 females); ketamine treated *Ift88* cHET, *n* = 6 (3 males + 3 females); ketamine treated *Ift88* cKO, *n* = 9 (5 males + 4 females). (**B**) Quantification of the fraction of successful pellet reach attempts/trial out of all reach attempts. Note significantly decreased motor learning rate in ketamine treated *Ift88* cKO group. (**C**) Line graph showing total number of pellet reach attempts for groups assayed in (**B**). Mice from all groups did not differ significantly in pellet reach attempts. Values in (**B**-**C**) represent mean ± standard error (SEM). \*\*\*\**P* < 0.0001, \*\**P* < 0.01. The statistical test used in (**B**-**C**) was a 2-way repeated measures ANOVA with post-hoc Tukey’s multiple comparisons test. Abbreviations: PRA – pellet reach assay, VEH – vehicle, KET – ketamine, PMC – primary motor cortex, CST – corticospinal tract, I.P.-intraperitoneal.

In correlation with the immunohistochemical data on caspase activation (Fig. 1), we observed a strongly reduced success rate at the pellet reach task solely in the ketamine-exposed *Ift88* cKO group (Fig. 3B). On the other hand, ketamine treated *Ift88* cHET animals and the vehicle groups exhibited no change in motor skill acquisition. This discrepancy in motor performance of the ketamine treated *Ift88* cKO mice could not be explained by lack of motivation in the ciliopathic animals, given that their mean total reaching attempts were significantly higher compared to cHET experimental groups at later stages (Fig. 3C). These data pointed to the possibility that elevated cleaved caspase 3 might be responsible for structural changes in deep layer neurons that lead to impairments in fine motor skill learning.

### *Ift88* cKO mice given ketamine have reduced apical spine density in layer V motor pyramidal neurons

Pyramidal neuron dendritic spine turnover has been closely associated with the process of memory consolidation, behavioral flexibility, and learning such as motor skill learning(*38, 39*). Indeed, spine content and plasticity in neocortical motor pyramidal neurons are directly correlated with rodent performance on the accelerated rotarod task, which is impaired in contexts of reduced spine plasticity such as during ageing(*40, 41*). Given that caspase 3 activity is known to influence the pruning of dendritic branches and spines in a non-apoptotic manner(*31*), we investigated spine density and dynamics of mature layer V motor pyramidal neurons in ketamine treated cohorts. To assess possible dendritic spine changes after ketamine-induced non-apoptotic caspase activation and the effect on motor skill learning, we performed chronic cranial window surgeries on ketamine treated control (*Ift88* cHET; *Thy1-GFP(M)*) and ciliopathic (*Ift88* cKO; *Thy1-GFP(M)*) mice, followed by multiphoton imaging and tracking of dendritic spines on the accelerated rotarod (Fig. 4A). The *Thy1-GFP(M)* transgenic reporter allows for sparse, yet bright labeling of individual dendritic segments and spines in apical arbors of neocortical layer V pyramidal neurons (Fig. 4B).

**Figure 4.**
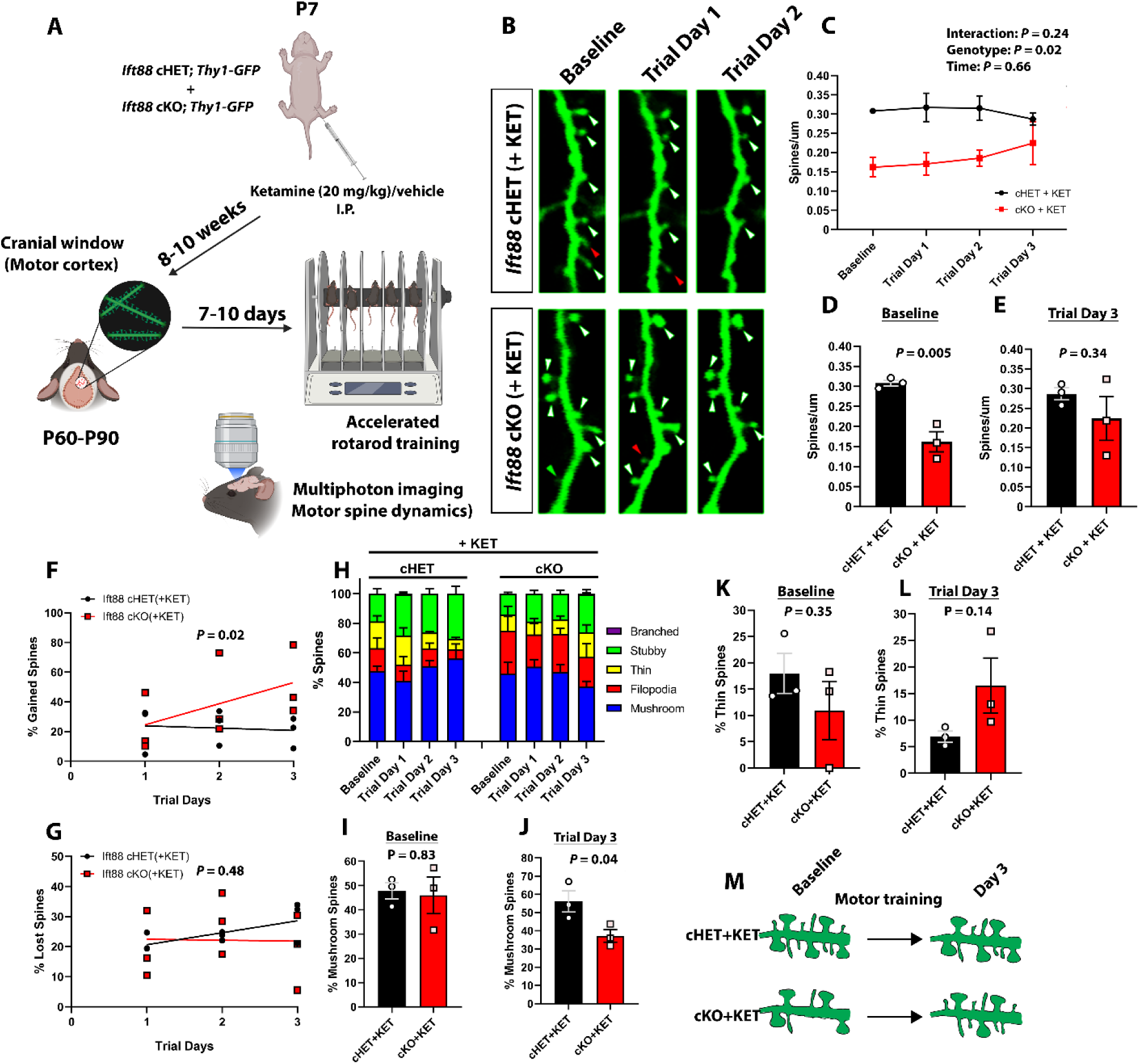
*Ift88* cKO mice given ketamine have reduced baseline layer V apical spine density in the motor cortex. (**A**) Diagram depicting experimental design and timeline for assessing effects of neonatal ketamine on apical dendritic spine dynamics of layer V pyramidal neurons in control (*Ift88* cHET) and forebrain-specific ciliopathy (*Ift88* cKO) mice containing the transgenic *Thy1- GFP(M)* reporter allele during motor training with the accelerated rotarod. n = 3 mice per group (2 males + 1 female in both groups). (**B**) Representative micrographs of layer V apical dendritic branch segments and spines tracked over the course of motor training with multiphoton imaging. White arrows show spines detected stably across several imaging sessions; red arrows indicate spines lost by the next imaging session; green arrows indicate spines gained by next imaging session. (**C**) Line graph showing quantification of overall dendritic spine density at baseline and following subsequent rotarod training sessions. Spine density was notably lower at baseline in ketamine treated *Ift88* cKO mice compared to controls and displayed a slow increase by the final training session. (**D-E**) Bar chart comparisons of apical spine density at baseline and final training days, respectively, between experimental groups. (**F-G**) Line graphs depicting the rates of dendritic spine gain or loss, respectively, during rotarod motor training in ketamine treated control (*Ift88* cHET) and forebrain-specific ciliopathy (*Ift88* cKO) groups. Note continuous gain of new spines in *Ift88* cKO mice throughout training, while control spine gain plateaus following first training session. Regression trendlines are shown for each group, with *P* values indicating statistical significance of slope comparisons. (**H**) Statistical diagram of percentages of classified dendritic spines in layer V pyramidal neurons imaged during the motor training. Spine class color mapping is shown next to the graph. (**I-J**) Bar charts displaying fraction of mushroom/mature dendritic spines at baseline and final training day, respectively. Note decrease detected in the *Ift88* cKO group at the end of training. (**K-L**) Charts showing fractions of thin/immature spines detected in both groups at baseline and final training day, respectively. Note trend towards increase in thin spines on final training day in *Ift88* cKO mice exposed to ketamine. (**M**) Schematic summarizing effects of motor training on layer V apical spines in control and *Ift88* cKO animals following perinatal ketamine administration. Values represent mean ± standard error (SEM). \**P* < 0.05, \*\**P* < 0.01. The statistical test used in **C** was a 2-way repeated measures ANOVA. Data in **F**-**G** was analyzed using regression analysis. Data in **D-E** and **I**-**L** was analyzed using unpaired student’s t tests. Abbreviations: KET – ketamine, I.P.-intraperitoneal.

Individual dendritic segments located in neocortical layer I were tracked during training, and their spine content and morphology were examined. Overall dendritic spine density was strongly reduced in the *Ift88* cKO group at baseline conditions, prior to the onset of accelerated rotarod training, and it displayed a slow increase up until the final training day (Fig. 4C-E). Control spine density remained at a higher steady level through most training days, displaying a subtle decline by the end of the experiment. This finding is consistent with the interpretation that a single dose of neonatal ketamine treatment reduces apical spine density in ciliopathic pyramidal neurons and leads to a dampened motor learning rate during the initial skill acquisition phase.

We next examined the dynamics of individual spines between imaging sessions, by quantifying the proportion of gained or lost spines relative to their baseline numbers (Fig. 4F-G).

The control group dendrites experienced a ∼23% gain in spines between baseline and first trial imaging sessions, which plateaued and declined by the final training day. In contrast, the ciliopathic dendrites matched the initial rate of spine gain yet continuously gained new spines until the end of motor training, displaying a mean ∼51% increase in gained spines compared to baseline levels (Fig. 4F). To examine the relationship between spine gain and time more closely we performed linear regression analysis on data obtained between trial days 1 and 3. The rate of spine gain displayed by *Ift88* cKO animals during this time window was significantly higher compared to controls, indicating enhanced spine addition in ciliopathic mice (Fig. 4F). In contrast, the rate of spine loss did not differ between the two conditions (Fig. 4G) indicating an overall net spine gain in ketamine-exposed ciliopathic layer V neurons during training. These results point to continuous addition of new spines during motor learning in the ciliopathic group leading up to the final rotarod training day, whereas control dendrites exhibit a more balanced spine turnover profile, favoring spine loss in the later training phases.

To understand whether motor training had differential impacts on different dendritic spine classes, individual spines were classified using morphological parameters, as described previously(*42*). Overall, we observed all major spine classes in both experimental groups across all imaging sessions, including mushroom, filopodial, thin and stubby spines (Fig. 4H). Branched spines were extremely rare, with only a single observation. Ciliopathic apical dendrites displayed a trend towards an inverted frequency profile of mushroom and thin spines compared to controls, with a tendency towards decrease or increase respectively, by the end of motor training. Mushroom spines, which represent mature postsynaptic sites forming stable synapses, did not differ between groups in their proportions at baseline (Fig. 4I). However, by the final training day their proportion decreased by ∼40% in ciliopathic mice (Fig. 4J). Thin spines, which are transient and associated with learning, exhibited greater variability within groups, and while baseline differences are unclear (Fig. 4K), their mean proportion displayed an increase (∼82%) in ciliopathic animals in the final imaging sessions (Fig. 4L). These findings indicate that while motor training impacts spine turnover in both groups, ciliopathic mice exhibit a spine profile characteristic of continuous learning which is protracted compared to controls, which acquire a more mature spine profile and diminish their spine plasticity by the end of training.

In summary, our spine dynamics profiling experiments show evidence of decreased baseline apical spine density in ciliopathic layer V pyramidal neurons exposed to ketamine (Fig. 4M). Consistent with a delay in fine motor skill learning (Fig. 3), ciliopathic neurons displayed protracted spine plasticity during motor training, with a delay in acquisition of mature postsynaptic sites by the final training session.

### Pharmacological inhibition of caspase signaling silences caspase activation in ciliopathic mice due to ketamine

To confirm the involvement of caspases in ketamine-induced protracted spine plasticity and motor learning impairments in ciliopathic mice, we utilized quinolyl-valyl-O-methylaspartyl-[-2,6- difluorophenoxy]-methyl ketone (QVD-Oph), a non-competitive inhibitor of caspases 1, 3, 8, and 9, which can cross the blood-brain barrier and displays no overt toxicity *in vivo*(*43*). To confirm effective inhibition of caspase 3 activation in our model, we administered two doses of QVD-Oph (10 mg/kg each, intraperitoneally) with the first dose given immediately following ketamine treatment, and the second dose 12 hours after the first (Fig. 5A). As in the initial experiments (Fig. 1), CC3 immunoreactivity was assessed in conjunction with NeuN in the deep layers of the PMC. Treatment with QVD-Oph completely suppressed neuronal CC3, bringing signal to below control (*Ift88* cHET^+^ ketamine) levels (∼70% reduction) (Fig. 5B-C). This reduction was reflected across different signal intensities with particularly lower CTCF values among more strongly immunoreactive neurons (Fig. 5D-E). While QVD-Oph treatment strongly reduced caspase 3 activation due to ketamine in the *Ift88* cKO motor cortex, it resulted in a more subtle reduction trend (∼16.5%) in Iba1^+^ microglial cells (Supp. Figure S3C), which was not statistically significant. These findings indicate that a short treatment regime with a pan-caspase inhibitor, administered promptly after ketamine exposure, is sufficient to rescue neuronal caspase 3 activation in the deep layers of the ciliopathic PMC.

**Figure 5.**
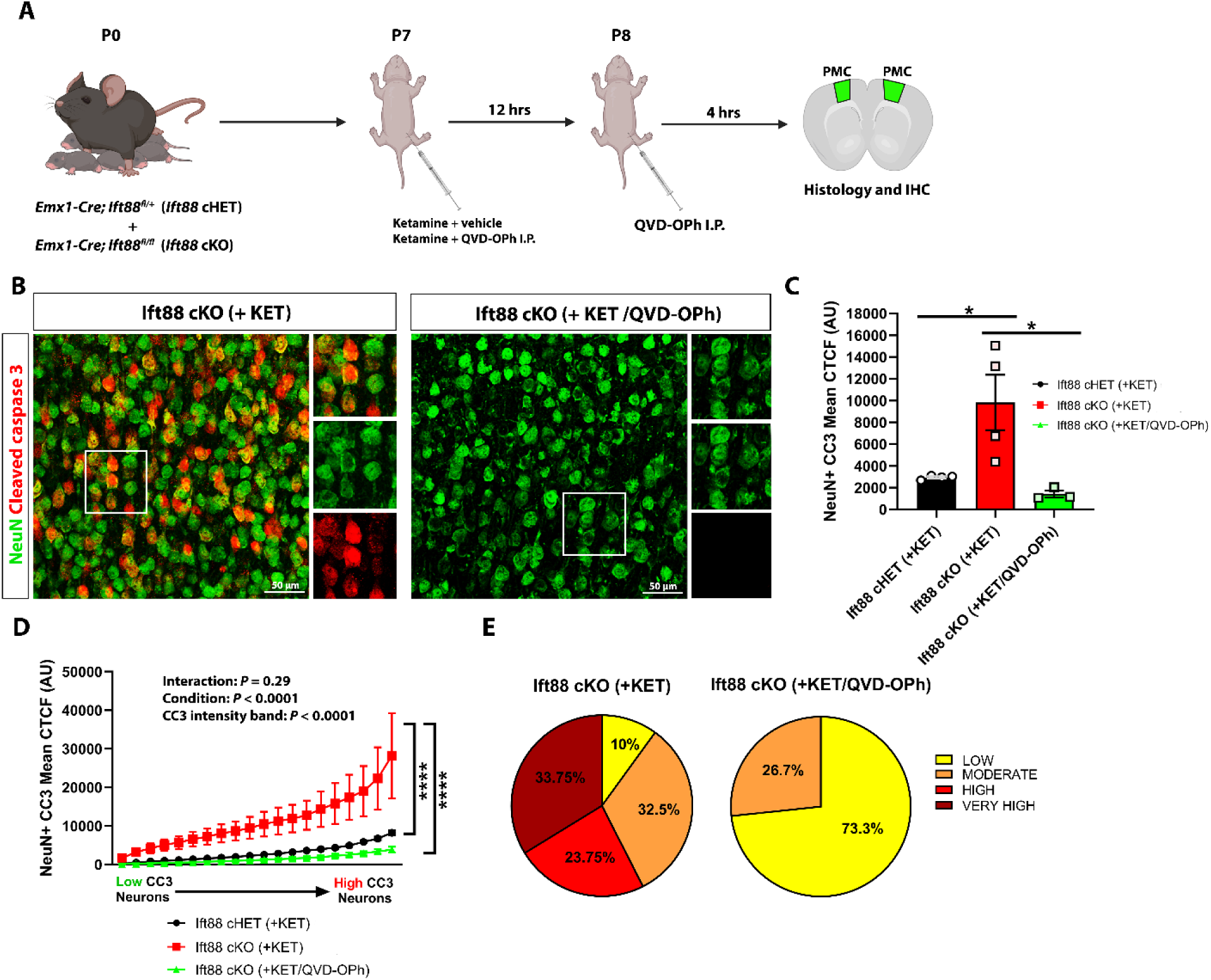
Perinatal treatment with pan-caspase inhibitor QVD-Oph suppresses neuronal caspase activation due to ketamine in *Ift88* cKO mice. (**A**) Schematic exhibiting ketamine and QVD-Oph treatment regime in perinatal *Ift88* cKO animals, followed by histological assessment of caspase 3 activation in neocortical neuronal cells. (**B**) Representative micrographs of primary motor cortex (PMC) layer V/VI showing magnified insets (white boxes) of cleaved caspase 3 (CC3) immunoreactivity co-localizing with neuronal NeuN in ketamine and ketamine + QVD- Oph treated *Ift88* cKO mice. (**C**) Quantification of the mean CC3 CTCF across sampled deep layer neurons from groups shown in **B** alongside control ketamine treated *Ift88* cHET group. (**D**) Line graph displaying mean cleaved caspase 3 (CC3) corrected total cell fluorescence (CTCF) values across 20 representative neurons/PMC region of interest (ROI) (ordered from neurons with lowest to neurons with highest mean CC3 CTCF), and their comparison between ketamine treated *Ift88* cHET/cKO and ketamine + QVD-Oph treated *Ift88* cKO mice. Note normalization of CC3 intensity in deep layer neurons of ketamine + QVD-Oph treated *Ift88* cKO mice. Values show means ± standard error (SEM). \**P* < 0.05,\*\*\*\**P* < 0.0001. The statistical test used in **C** was a 1- way ANOVA with post-hoc Tukey’s multiple comparisons test (n=3-4 each, 2 males + 2 females in cHET/cKO groups treated with ketamine and 2 males + 1 female in cKO group treated with ketamine/QVD-Oph). Data in **D** was analyzed using a 2-way ANOVA with post-hoc Tukey’s multiple comparisons test (n=3-4 in each). (**E**) Pie charts showing proportions of neurons displaying low (0-2000), moderate (2000-6000), high (6000-10,000) and very high (10,000+) CC3 CTCF values in ketamine and ketamine + QVD-Oph groups (n=3-4 each). Abbreviations: CC3- cleaved caspase 3, KET-ketamine, I.P.-intraperitoneal.

### Caspase suppression rescues fine motor deficits and apical spine density in *Ift88* cKO motor cortex

To investigate the consequences of dampened caspase 3 activation on fine motor skills and dendritic spines of motor layer V pyramidal neuron apical dendrites, we employed the same experimental paradigm used during neonatal ketamine exposure (Fig. 6A). One set of cohorts was fasted for pellet reach fine motor skill acquisition, while the other underwent chronic cranial window implantation over the motor cortex, followed by accelerated rotarod training and concurrent spine multiphoton imaging (Fig. 6A).

**Figure 6.**
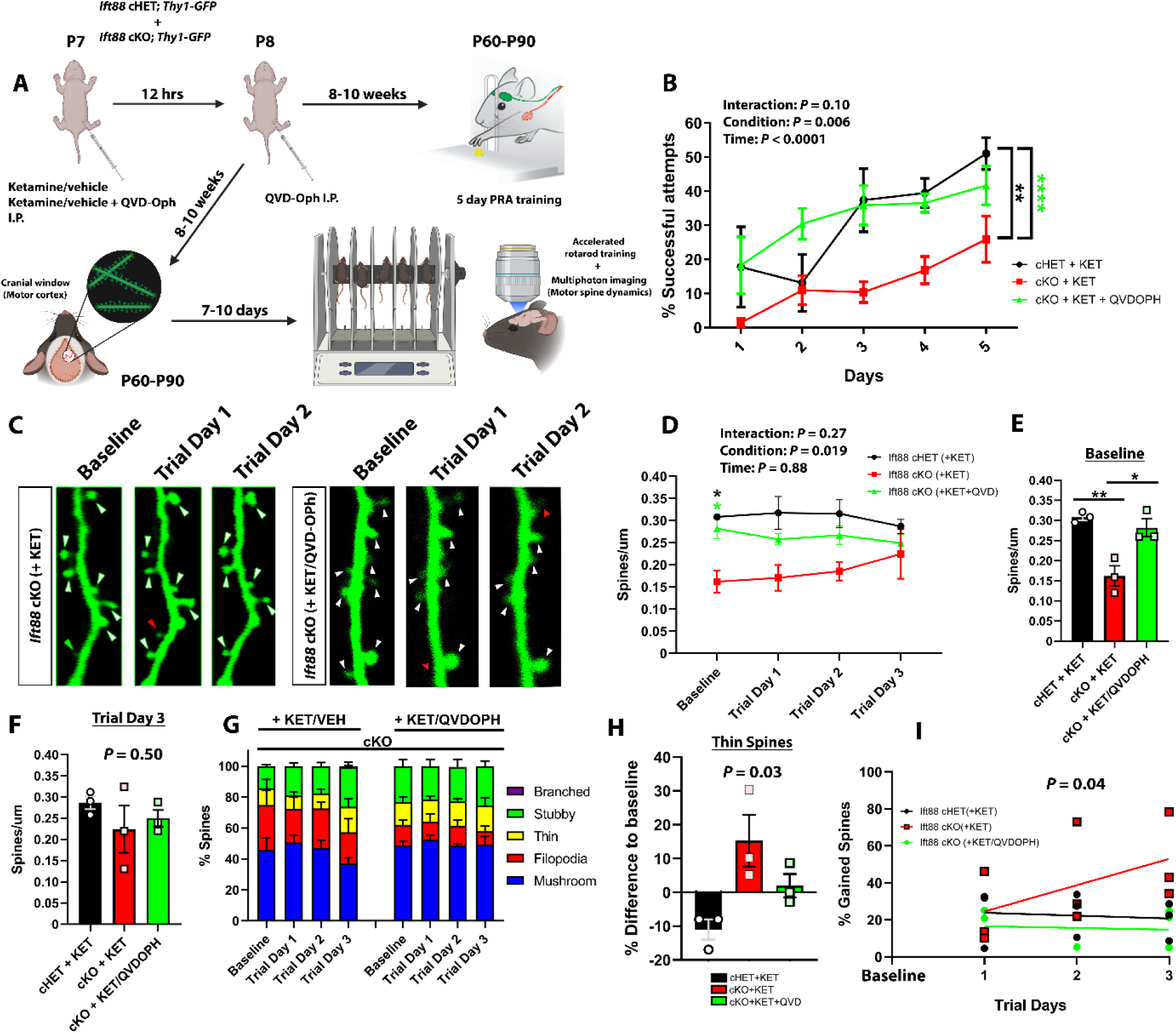
Perinatal caspase signaling inhibition rescues fine motor skill and apical motor spine density due to ketamine in *Ift88* cKO mice. (**A**) Schematic depicting ketamine and QVD-Oph treatment regime, followed by assessment of pellet reach fine motor skills or dendritic spine dynamics of layer V pyramidal neurons in ketamine and ketamine + QVD-Oph Ift88 cKO animals. (**B**) Quantification of the fraction of successful pellet reach attempts/trial in ketamine treated groups and ketamine + QVD-Oph treated ciliopathic (*Ift88* cKO) mice. Note significantly improved and rescued motor learning rate in ketamine + QVD-Oph treated *Ift88* cKO group. Numbers used: ketamine treated *Ift88* cHET, *n* = 6 (3 males + 3 females); ketamine treated *Ift88* cKO, *n* = 9 (5 males + 4 females); ketamine + QVD-Oph treated *Ift88* cKO, *n*=5 (3 males + 2 females). (**C**) Representative micrographs of layer V apical dendritic branch segments and spines tracked over the course of motor training with multiphoton imaging in ketamine and ketamine + QVD-Oph treated *Ift88* cKO groups (*n* = 3; 2 males + 1 female in all groups). White arrows show spines detected stably across several imaging sessions; red arrows indicate spines lost by the next imaging session; green arrows indicate spines gained by next imaging session. (**D**) Line graph showing quantification of overall dendritic spine density at baseline and following subsequent rotarod training sessions. Spine density was enhanced at baseline in ketamine + QVD-Oph treated *Ift88* cKO mice compared to group receiving only ketamine. (**E-F**) Bar chart comparisons of apical spine density at baseline and final training days, respectively, between experimental groups. The statistical tests used were 1-way ANOVA with post-hoc Tukey’s multiple comparison tests. (**G**) Statistical diagram of percentages of classified dendritic spines in layer V pyramidal neurons imaged during the motor training. Spine class color mapping is shown next to the graph. (**H**) Quantification of the change in thin/immature spines relative to baseline during motor learning. Note opposing trends in *Ift88* cHETs and cKOs, with QVD-Oph treatment stabilizing thin spine turnover. (**I**) Line graph depicting the rates of dendritic spine gain during rotarod motor training in ketamine and ketamine + QVD-Oph treated ciliopathic (*Ift88* cKO) groups. Ketamine treated *Ift88* cHET data is shown as a control. Note trend towards diminished gain of new spines in ketamine + QVD-Oph treated *Ift88* cKO mice following the first training day, mimicking the control group spine gain plateau. Rates of spine loss were similar at the end of motor training in all tested groups. Values represent mean ± standard error (SEM). \**P* < 0.05; \*\**P* < 0.01. The statistical test used in **B** and **D** was a 2-way repeated measures ANOVA with post-hoc Tukey’s multiple comparisons test. The data in **E**-**F** and **H** was analyzed using a 1-way ANOVA with post-hoc Tukey’s multiple comparisons test. Data in **I** was analyzed using regression analysis. Abbreviations: KET-ketamine, I.P.-intraperitoneal.

Brief QVD-Oph treatment given following ketamine-induced acute caspase activation resulted in the complete rescue of fine motor skill acquisition on the pellet reach task, represented by the enhanced success rate of pellet acquisition in *Ift88* cKO mice that matched controls (ketamine-treated *Ift88* cHETs) (Fig. 6B). This finding strongly argues that the initial upregulation in neuronal caspase 3 activation following ketamine exposure is critical for the fine motor skill deficit seen in ciliopathic *Ift88* cKO animals.

Given that the observed fine motor skill acquisition delay in ketamine treated *Ift88* cKOs was associated with altered apical spine density and turnover in layer V pyramidal neurons of the PMC, we assessed QVD-Oph treated spines during motor learning (Fig. 6C). Quantification of overall spine density revealed a complete rescue in QVD-Oph treated mice at baseline, and a return to control levels (Fig. 6D-E). After the final training day, spine density did not differ significantly between the experimental groups (Fig. 6F). This result indicates that suppressing caspase 3 activation following neonatal ketamine sedation is sufficient to restore apical spine density to control levels, and that exacerbated caspase signaling mediates apical spine pruning with resultant density deficits that persist into adulthood. Spine classification at baseline and during motor learning trials revealed a tendency towards reduction in filopodial spines in QVD-Oph treated mice across all imaging sessions compared to ketamine exposed *Ift88* cKOs (Fig. 6G; Supplementary Fig. S3C). This reduction resulted in overall levels of filopodial spines that mimicked controls (ketamine treated *Ift88* cHETs). Mushroom and thin spine fractions also exhibited a propensity towards a return to control levels at final trial day and baseline respectively (Supplementary Fig. S3D-G). Thin immature spines exhibited an increase in proportion at the final trial day relative to baseline in ciliopathic mice, which was significantly different to the relative decrease seen in controls (Fig. 6H). QVD-Oph normalized this elevation in thin spines, although this was not statistically significant. However, the thin spine fraction in QVD-Oph treated animals remained at higher overall levels than controls in the final imaging session (Supplementary Fig. S3F), arguing that this component of the spine phenotype is more dependent on the ciliopathy than ketamine- induced caspase 3 activation.

To understand whether QVD-Oph treatment alters apical spine turnover during motor learning, we quantified the fraction of gained and lost spines as before. QVD-Oph restored the spine gain profile in *Ift88* cKO animals to that seen in controls, likely due to the normalization of baseline spine density (Fig. 6I). Linear regression analysis found no statistically significant difference in the rate of spine loss between all experimental groups (Supplementary Fig. S3G). Overall, our findings indicate that caspase 3 activation, due to acute neonatal ketamine exposure in ciliopathic animals, is a key driver of reduced apical spine density and altered spine turnover in layer V pyramidal neurons of the adult PMC. Brief pharmacological inhibition of caspase signaling following ketamine administration is capable of completely restoring apical spine density and fine motor skill acquisition to control levels, while stabilizing spine turnover rates during motor learning.

## Discussion

Our study here demonstrates for the first time that neonatal exposure to a single dose of ketamine is capable of inducing caspase 3 with sub-lethal yet detrimental consequences on neuronal structure and spine plasticity in ciliopathic pyramidal neurons, leading to motor learning impairments. While a complete loss of primary cilia is not frequently encountered in patient populations, apart from rare genetic conditions such as primary ciliary dyskinesia(*44*), genetic variants with damaging effects on cilia integrity and function are frequent in patients with CHD(*3, 45, 46*). Anesthesia and sedation are currently critical components of care in infants with CHD for various surgical and/or interventional procedures(*47, 48*). Given the increased frequency of motor skill impairments in individuals with CHD and the high prevalence of mutations associated with ciliary dysfunction, our findings suggest potentially harmful interactions between anesthetic exposures and genomic variants associated with primary cilia deficits in the CHD population. In addition, our results imply a capacity for acute ketamine sedation to induce caspase-dependent, non-apoptotic neuronal structural changes long-term, which to our knowledge has been underappreciated in the field. Notably, we demonstrate that brief treatment with the pan-caspase inhibitor QVD-Oph, known to possess low *in vivo* toxicity, rescues the delay in motor learning and deficits in neuronal structure and spine plasticity, revealing brief inhibition of caspases as a potential therapeutic target for children with CHD who require multiple sedation procedures during the early postnatal period.

Prior work by Ishii et al(*27*) demonstrated that perinatal ethanol exposure causes caspase-induced degenerative changes to neuronal arbors in ciliopathic *Ift88* cKO mice, which occur due to downregulation of the Akt pathway. Pharmacological activation of insulin-like growth factor 1 (IGF1)-Akt signaling was shown to restore dendritic arborization in the ciliopathic model. Interestingly, caspase 3 activation was present after ethanol treatment in neurons where ciliary loss was induced from P5, suggesting the adverse interaction occurs primarily within the perinatal window, rather than from earlier embryonic loss of cilia such as occurs with the *Emx1-Cre* driver. Activation of the IGF1 receptor was localized at the base of the primary cilium, near the ciliary transition zone, a diffusion barrier controlling the trafficking of ciliary proteins, which is particularly enriched for damaging de novo variants in cases of CHD(*4, 49*). Our study extends these results to ketamine, another NMDA receptor antagonist, with likely shared mechanistic features with ethanol exposure in a ciliopathic context.

Complementary to the ethanol findings, we detected a specific subpopulation of layer V neurons susceptible to ketamine-induced caspase 3 activation, pointing to possible differences in susceptibility to environmental agents among neighboring cells. Neuronal degeneration and death are often accompanied by neuroinflammatory infiltration by astrocytic and microglial cells(*50*); however, we did not detect robust changes in microglia following ketamine administration in ciliopathic mice. One possible explanation is that the sampled time point (16 hours post-treatment) is too early to detect the full extent of microglial expansion. Alternatively, the level of neuronal caspase 3 activation in our model does not reach a critical threshold which would trigger robust neuroinflammatory changes in the PMC. Indeed, the observed augmentation of caspase signaling might be elevated enough to trigger cytoskeletal fragmentation and pruning of dendrites and spines, yet not so high to cause neuronal death. Whether inhibitor of apoptosis (IAP) proteins specifically limit the progression to neuronal apoptosis in the ciliopathic context is unknown and would benefit from future study.

Our motor skill behavioral analysis detected robust impairments of fine motor skill learning in the ciliopathic animal group following a single dose of ketamine. Since we did not observe evidence of reduced motivation in ketamine treated *Ift88* cKOs, it is plausible that either components of motor initiation or integration are impaired in the ciliopathic mice. Curiously, visuomotor integration is a particularly vulnerable behavioral domain in patients with CHD(*51, 52*), which could be at risk due to adverse interactions between anesthetics/sedatives and susceptible genetic backgrounds. Concurrently with the fine motor skill impairments, ciliopathic mice treated with ketamine showed a striking reduction in apical spine density and protracted spine turnover profiles, structural correlates of reduced synaptic plasticity and delayed learning. Interestingly, the spine density deficit was confined to the apical rather than basal compartment, although this might be explained by the fact that we assessed basal dendritic spines at the end of motor training, rather than throughout learning. Apical spine class assessments showed a tendency towards fewer mature spines and a greater fraction of immature spines by the end of motor training in ciliopathic mice after ketamine, indicating delayed plasticity and consequently motor learning. Repeated exposures to ketamine between the second and third postnatal week have previously been shown to dampen spine plasticity in adulthood(*53*). Given the observed caspase-dependent spine density impairment at the onset of motor training, the likely explanation for the protracted spine turnover profile is the propensity for adding new immature spines coupled with a paucity of mature, motor memory-consolidating spines. Whether the spine density defect occurred primarily due to initial caspase-dependent pruning(*54*) following ketamine or was also shaped secondarily by altered spinogenesis and spine dynamics prior to training(*55*) is not clear and would be an interesting avenue to investigate further.

In addition to the spine deficits, we found evidence of a loss of dendritic complexity in ciliopathic pyramidal neurons after a single ketamine exposure, which coupled with the spine impairment points to substantial caspase-driven morphological changes resulting due to an interaction between ciliary dysfunction and ketamine exposure. Prior studies examining the anti- depressant effects of ketamine in wild-type rats showed that acute ketamine treatment increased short-term prefrontal cortical neuron dendritic arborization and spine density, likely via enhanced mammalian target of rapamycin (mTOR) signaling(*56, 57*). These discrepancies could reflect differing time windows of ketamine exposure, as well as the fact that our study investigated the interaction with a loss of primary cilia. In contrast with previous work which focused more on the classical caspase-driven apoptosis pathway, our study suggests non-apoptotic, long term cytomorphological changes could be a cause for concern in patient populations receiving anesthesia that are genetically predisposed to ciliary impairments or carry IGF1-Akt pathway polymorphisms(*58*), such as in CHD(*5, 59, 60*).

Given that caspases are ubiquitous housekeeping enzymes during development, long-term caspase inhibition potentially has off-target side effects. Thus, we tested a potent pan-caspase inhibitor, QVD-Oph, during a limited period that specifically targets caspase activation during/immediately after neonatal ketamine administration. Our results indicate that a brief exposure to a pan-caspase inhibitor is sufficient to correct the motor behavior impairments and spine deficits induced by ketamine. QVD-Oph use has not been associated with toxicity in vivo and has shown efficacy in limiting caspase activation in models of stroke, spinal cord injury and neurodegenerative disorders(*43*). In our model the inhibitor rescued both the fine motor skill learning rate and the baseline apical spine density, pointing to caspase-dependent pruning as the likely cause of the behavioral impairment. Caspase inhibition also restored the fraction of mature motor spines, possibly aiding the process of earlier memory consolidation compared to ciliopathic mice exposed to neonatal anesthesia. Given the safety profile of QVD-Oph and its efficacy in inhibiting both initiator and effector caspases with brief treatment, it shows promise for limiting caspase-dependent neuronal structural impairments due to agents such as sedatives and general anesthetics.

While this work identifies a clear interaction between ketamine and loss of primary cilia in the neonatal neocortex, there are several limitations. Firstly, complete ablation of primary cilia in a systemic manner is incompatible with life, and severe ciliopathies such as Meckel-Gruber syndrome lead to perinatal lethality(*61*). The ciliopathic mouse model allows for targeted loss of primary cilia in excitatory neurons of the neocortex from late mid-gestation, providing the means to test cell-specific sensitivity to the interaction between ketamine and ciliary loss. Partial ciliary loss, primary cilia shortening and/or aberrant function are more frequently encountered among patients, and future modelling efforts taking this into account would help resolve whether this detrimental interaction occurs in more common clinical scenarios. Secondly, given that ketamine signals through many of the same pathways as most inhalable anesthetics, determining whether similar interactions occur between agents such as sevoflurane and primary cilia impairments would be highly pertinent. In addition, neonatal anesthesia for major surgery such as neonatal CHD repair typically involves a mix of compounds including benzodiazepines, propofol, sevoflurane combined with fentanyl and dexmedetomidine, which in combination might result in caspase-induced apoptosis and cell death, rather than sublethal structural remodeling as we have observed. Testing the interaction between these compounds and damaging ciliary variants would be feasible in the mouse and would yield information on compound-specific effects on neuronal morphology and motor function. The ketamine dose, while relatively low, points to heightened sensitivity of ciliopathic animals to ketamine-induced sedation, which would likely be exacerbated by higher, continuously administered doses encountered clinically. Thirdly, exploring the mechanistic basis of the sublethal caspase effects on neuronal spines, and how this signaling cascade is triggered by ketamine exposure is necessary in future studies to understand how caspase inhibition might mediate its neuroprotective effects on neuronal structure and function.

To conclude, our study demonstrates that even a single exposure to ketamine sedation can induce detrimental, caspase-driven morphological changes to motor pyramidal neurons as well as motor learning deficits in ciliopathic subjects. The caspase-mediated effects are non-apoptotic and reversible through early and brief caspase inhibition.

## Materials and Methods

### Animals and treatments

*Ift88^fl/fl^* (B6.129P2-*Ift88^tm1Bky^*/J) transgenic mice on a C57BL6/J background were used in this study and were obtained from Jackson Laboratory (Strain #:022409; RRID:IMSR_JAX:022409). The *Ift88^fl/fl^*line was initially crossed to *Emx1-Cre* (B6.129S2-*Emx1^tm1(cre)Krj^*/J; Strain #:005628; RRID:IMSR_JAX:005628) to generate conditional heterozygous (*Ift88* cHET) or homozygous knockout (*Ift88* cKO) mice. The resulting animals were then crossed to the *Thy1-GFP(M)* line (STOCK Tg(Thy1-EGFP)MJrs/J; Strain #:007788; RRID:IMSR_JAX:007788) to generate *Emx1- Cre; Ift88^fl/+^;Thy1-GFP* (*Ift88* cHET; *Thy1-GFP*) or *Emx1-Cre; Ift88^fl/fl^;Thy1-GFP* (*Ift88* cKO; *Thy1-GFP*) animals for neuronal arborization and dendritic spine imaging. All animals were housed on a 12-h light/dark cycle and, unless required to be different by the experiment, provided free access to a standard rodent food pellet diet and water.

The treatment regime consisted of administering a single dose of ketamine (20 mg/kg), dissolved in 0.9% normal saline (vehicle), or vehicle intraperitoneally to postnatal day seven (P7) pups born to *Emx1-Cre; Ift88^fl/+^;(Thy1-GFP) x Ift88^fl/f^;(Thy1-GFP)* crosses. For the caspase inhibitor experiments, a single dose of ketamine (20 mg/kg) was followed up by two doses of QVD-Oph (10 mg/kg) dissolved in 1% DMSO in 0.9% normal saline. The first QVD-Oph dose was given right after ketamine, with the second dose administered at 12-13 hours following the first.

### Immunohistochemistry

To prepare histological samples for immunohistochemistry, animals at the indicated stages were intracardially perfused with ice cold phosphate buffered saline (PBS), followed by ice cold 4% paraformaldehyde (PFA) dissolved in PBS. Perfused brains were isolated and post-fixed in 4% PFA solution overnight at 4 °C. Following fixation, brains were sectioned coronally into 30 or 300 µm slices using a vibratome (Leica VT1000 S) or placed into graded sucrose solutions (15% then 30% sucrose in PBS) prior to freezing for cryosectioning into 20 µm coronal sections using a cryostat. Vibratome slices were used for free floating immunohistochemical staining for neuronal nuclear antigen (NeuN, EMD Milipore #ABN90)), cleaved caspase 3 (CC3, Cell Signaling Technologies #D175), ionized calcium binding adaptor molecule 1 (Iba1, EMD Milipore #MABN92) or green fluorescent protein (GFP, Abcam #ab13970). Secondary antibodies used were Alexa Fluor conjugated. Cryosections were used for slide-mounted immunohistochemistry for NeuN and terminal deoxynucleotidyl transferase dUTP nick end labeling (TUNEL Assay Kit, Cell Signaling Technologies **#**64936). The free floating immunohistochemical procedure consisted of incubating the vibratome slices in donkey serum blocking solution (donkey serum in 0.2% Triton-X in 0.01M PBS) for 1 hour at room temperature, followed by primary antibody solutions overnight (all diluted 1:500) at 4 °C. On the following day, slices were rinsed three times for 5 minutes each with 0.01M PBS, after which they were incubated in secondary antibody solution for 2 hours at room temperature. This was followed by a final three rinses for 5 minutes each with 0.01M PBS before counterstaining with Hoechst (diluted 1:1000 in PBS, Invitrogen #33342 trihydrochloride-trihydrate) and mounting in Prolong anti-fade medium (Thermofisher #P36934). Primary and secondary antibodies were diluted in donkey serum blocking solution, apart from staining involving cleaved caspase 3 antibodies, which were diluted in Immunostain Enhancer (Pierce #46644) for signal amplification.

### Image analysis and quantification

Confocal images of NeuN/CC3 and Iba1 staining were obtained using a Nikon A-1 HD25 confocal microscope. GFP staining from Thy1-GFP neuronal and dendritic spine samples was imaged using a Leica SP8 confocal system. High resolution (1024 x 1024 pixels) z series scans with a 1 μm z step interval and consisting of 25-30 images were taken for NeuN/CC3 and Iba1 stained slides with a 20x air objective and 3x optical zoom. Two samples of the region of interest (ROI) (one from either hemisphere) from 3 slices/animal were acquired from the primary motor cortex (PMC) deep layers (layers V-VI) for downstream analysis. The sampled coronal slices were taken from brain regions equivalent to stereotaxic coordinates +1.0mm, +1.5mm and +2mm [anteroposterior], +/-1.5mm [mediolateral] and -1.0mm [dorsoventral] relative to bregma as defined in the Allen Brain Reference Coronal Atlas for adult mice. The xy dimensions of the imaging fields of view (∼300 x 300 μm) ensured that the acquired deep (layers V-VI) cortical layer ROIs were acquired in a non-overlapping manner with superficial (layers I-III) territories and were representative of PMC deep layers. The z series scans were initially flattened into single images using a maximum intensity z projection in the NIS-Elements software (Nikon). NeuN+ cells were inspected for CC3 staining, followed by marking the borders of the CC3 signal and quantification of the mean CC3 signal intensity and surface area from each NeuN+ cell. Mean background intensity was also obtained from areas surrounding the cells. To obtain corrected total cell fluorescence (CTCF) values for the CC3 signals, we used the formulas:

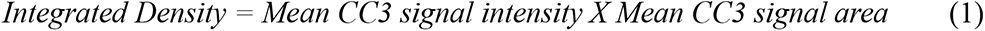

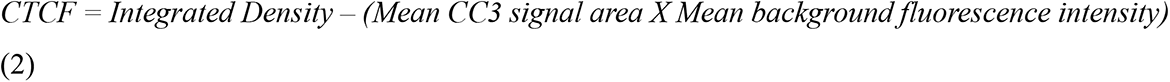

20 neurons were analyzed in this manner from each ROI (for a total of ∼80-120 neurons/animal), prior to averaging for comparisons between groups

For GFP stained slides, coronal slices (300 µm) from two independent section levels anterior to the bregma were imaged and individual neurons were used for morphological reconstruction and analysis with Neurolucida 360 software. High resolution (1024 x 1024 pixels) z series scans were obtained using either a 0.5 μm or 1 μm step interval for basal dendritic spines and neuronal arbors respectively. Confocal z stacks consisted of 40-60 or 100-150 images for spine and neuronal arbor imaging respectively. To maximize the number of basal spines and dendrites captured for morphometric analyses, tile scans were performed with 4 x 4, or 5 x 5 z tiles stitched together using 10% overlap with the Leica LAX software. Dendritic spines were imaged using an oil immersion 100x objective and a 2.5x optical zoom, while neuronal arbors were captured using a 20x air objective and 2.5x optical zoom. Tile z scans were imported into Neurolucida 360 (MBF) for three-dimensional neuronal arbor and dendritic spine tracing, and spine classification using default parameters. Once completed, individual traces were analyzed using Neurolucida Explorer (MBF) to obtain measures of branching complexity, soma volume, spine density, spine volume and spine class frequency.

### Cranial window surgeries

Following treatment at P7, transgenic mice (*Ift88* cHET; *Thy1-GFP* and *Ift88* cKO; *Thy1-GFP*) underwent chronic cranial window implantation surgeries at 2-3 months of age, as previously described(*62*), with slight modifications. Briefly, animals were anesthetized using a cocktail of ketamine/xylazine (100 mg/kg ketamine, 16 mg/kg xylazine, administered intraperitoneally) and placed on a body warmer pad (Kent Scientific #RT-0520) kept at 37 °C. The toe pinch reflex and breathing pattern were used to monitor the depth of anesthesia. Carprofen (5 mg/kg) and dexamethasone (2.5 mg/kg) were administered subcutaneously to minimize inflammation and brain edema. The scalp was shaved using a trimmer and eye ointment applied to prevent corneal drying. The animal was mounted on the stereotaxic instrument (Kopf Model #902) and 1% lidocaine was applied subcutaneously under the scalp. A single vertical incision was made over the skull, and the skin and connective tissue were retracted to expose the frontal skull bone. A 3-3.5 mm diameter circular craniotomy was performed over the motor cortex (craniotomy center at ∼1 mm anterior and 1 mm lateral to bregma), using a micro drill and regularly cooling the drill site with ice cold 0.9% saline solution. Absorbable hemostatic gelatin sponges (Fisher Scientific #NC0654350) dipped in ice cold 0.9% saline were used to stop and clear up any minor bleeding during the procedure. Following careful removal of the skull bone flap, a custom circular coverslip assembly (5 mm outer and 3 mm inner diameter) was used to replace the bone with a glass “window” as previously described(*62*). The glass window was glued to the surrounding bone using an instant adhesive (Loctite Super Glue liquid). A custom-built metal headpiece (Xometry) was then glued to the surrounding bone carefully to avoid contacting the glass window. Dental cement (Lang #1334CLR) was applied to the edges of the craniotomy and exposed areas surrounding the headpiece. Following this, the surgical animals were kept on the warming pad until they regained mobility.

### Multiphoton spine imaging

Surgical mice were allowed at least one week to recover from the procedure before imaging. To train animals on a motor task, we employed the accelerated rotarod. The mice were initially acclimated to the rotarod apparatus by placing them on a fixed speed (5 revolutions/min) regime for 15 minutes on the first day of testing. The mice were then trained on an accelerated rotarod for the next four days (5-40 revolutions/min over 5 minutes), for three trials/day, with trials separated by 5 minutes. All training trials were video recorded using a Basler GigE camera mounted on a tripod and Noldus EthoVision XT software. Trials were performed at the same time of the day to limit behavioral variations due to circadian rhythm fluctuations. Video recordings were scored by a blinded investigator. The rotarod acclimation and training sessions (apart from first training session) were followed up (∼2 hours) by daily imaging and tracking of individual apical dendritic spines in neocortical layer I, using a multiphoton microscope (Evident FVMPE-RS). Briefly, the mouse was anesthetized using a cocktail of ketamine/xylazine (2/3 of surgical dose) and placed on a warming pad kept at 37 °C. Head fixation was achieved by bolting the metal headpiece to custom-built holders printed three-dimensionally in ABS polymer (Xometry). Imaging was performed using a 25x water immersion multiphoton objective (Evident XLPLN25XWMP2) and a 3x or 10x optical zoom. To capture GFP fluorescence a Ti:sapphire Mai Tai femtosecond laser (Spectra Physics) was tuned to 950 nm for excitation, and the emitted light was collected with a high sensitivity GaAsP detector. A z series scan of a larger field of view was initially obtained at 3x optical zoom for an overview of the area of interest and storing of coordinates for repetitive imaging. Higher resolution (800 x 800 pixels) z scans of groups of dendritic spines were acquired at 10x optical zoom using a 0.5 μm z step interval and 50-100 images for each region of interest (ROI). Five ROIs were acquired and tracked for each animal in each imaging session (for a total of four imaging sessions/animal). Image acquisition was performed using the Fluoview software (Evident). The resultant z series images were analyzed using Fiji/ImageJ to obtain apical spine density, spine class frequency and to track spine turnover (total of ∼550 spines) during motor training. Spine classification was performed using spine diameter, length and length/diameter ratio parameters as described previously(*42*).

### Pellet reach assays

To assess fine motor skill acquisition, we employed the pellet reach assay in *Emx1-Cre; Ift88^fl/+^* (*Ift88* cHET) and *Emx1-Cre; Ift88^fl/fl^* (*Ift88* cKO) mice. Mice are capable of rapidly learning the task, involving forelimb extension to reach a food pellet, pronated grasping of the pellet and finally forelimb retraction until the pellet is consumed. Deficits in successful performance of the task have been associated with impairments of the motor cortex due to stroke(*63*), or experimental manipulation of motor cortical circuit activity through chemo- and optogenetic means(*64*). To motivate pellet reaching behavior, mice were initially fasted for 5 days by restricting food to elicit a 10-15% body weight loss, which was maintained during training trials. Following the fasting period individual mice were placed in a Plexiglass training chamber with a single vertical slit located on the front wall. Individual food pellets were placed in front of the slit opening at a distance which requires the animals to use their forelimb to reach the pellet. A successful reaching attempt was termed successful if the mouse could reach, grasp the pellet, and consume it. Missed or dropped pellets were labelled as unsuccessful reaching attempts. Individual mice were assayed for five days, with each daily trial lasting for 20 minutes or until the animal successfully reached 20 pellets, whichever came first. All training sessions were performed at the same time of the day to limit behavioral variations due to circadian rhythm fluctuations and were video recorded using a Basler GigE camera mounted on a tripod and Noldus EthoVision XT software. Trial recordings were scored by a blinded investigator.

### Statistical analysis

All statistical analyses were performed in GraphPad Prism 8. All data are represented as mean ± standard error (SEM) from at least three biological replicates for each experiment. Cleaved caspase 3 intensity and spine density comparisons between three or more groups were analyzed by one-way ANOVA. Two factor data such as Sholl analyses were analyzed with two-way ANOVA followed by post-hoc Tukey’s or Sidak’s tests. Comparisons between two independent groups were made using unpaired t-tests or non-parametric Kruskal-Wallis tests. Spine dynamics and motor learning performance data were analyzed using repeated measures two-way ANOVA tests followed by post-hoc Tukey’s multiple comparisons tests. Differences between groups were judged to be statistically significant when *p* < 0.05. Asterisks denote statistical significance p < 0.05 (*); p < 0.01 (**); p < 0.001 (***); p < 0.0001 (****).

## Ethics statement

All animal experiments were carried out in accordance with the NIH Guide for the Care and Use of Laboratory Animals and were approved by the Institutional Animal Care and Use Committee of Children’s National Hospital.

## Data availability

The datasets used and/or analyzed during the current study are available from the corresponding author upon reasonable request. Further enquiries can be directed to the corresponding author.

## Acknowledgements

We would like to thank Dr Nathan Smith for his assistance with training and surgical techniques. We would also like to acknowledge Dr Li Wang from the Neurobehavioral core and the Microscopy cores at the Children’s National Research Institute for the use of their equipment in this project. The authors are thankful for the generosity of the Foglia and Hill families who supported our studies. The graphic images in the manuscript were created with BioRender.com.

## Contributions

N.S. and N.I. designed the study and wrote the manuscript. N.S., Z.A., C.F.S., N.R., G.B., C.T. and M.T.R. analyzed the data. N.S. conducted the treatment, behavioral and in vivo imaging experiments. Z.A. conducted the histological processing experiments and optimized immunohistochemical protocols. K.H.T. provided the *Ift88* flox mouse line, revised and approved the final manuscript. V.J.T. and T.H. revised and approved the final manuscript.

