## Supplementary Data for "Ciliopathy interacts with neonatal anesthesia to cause non-apoptotic caspase-mediated motor deficits"

**Supplementary material**

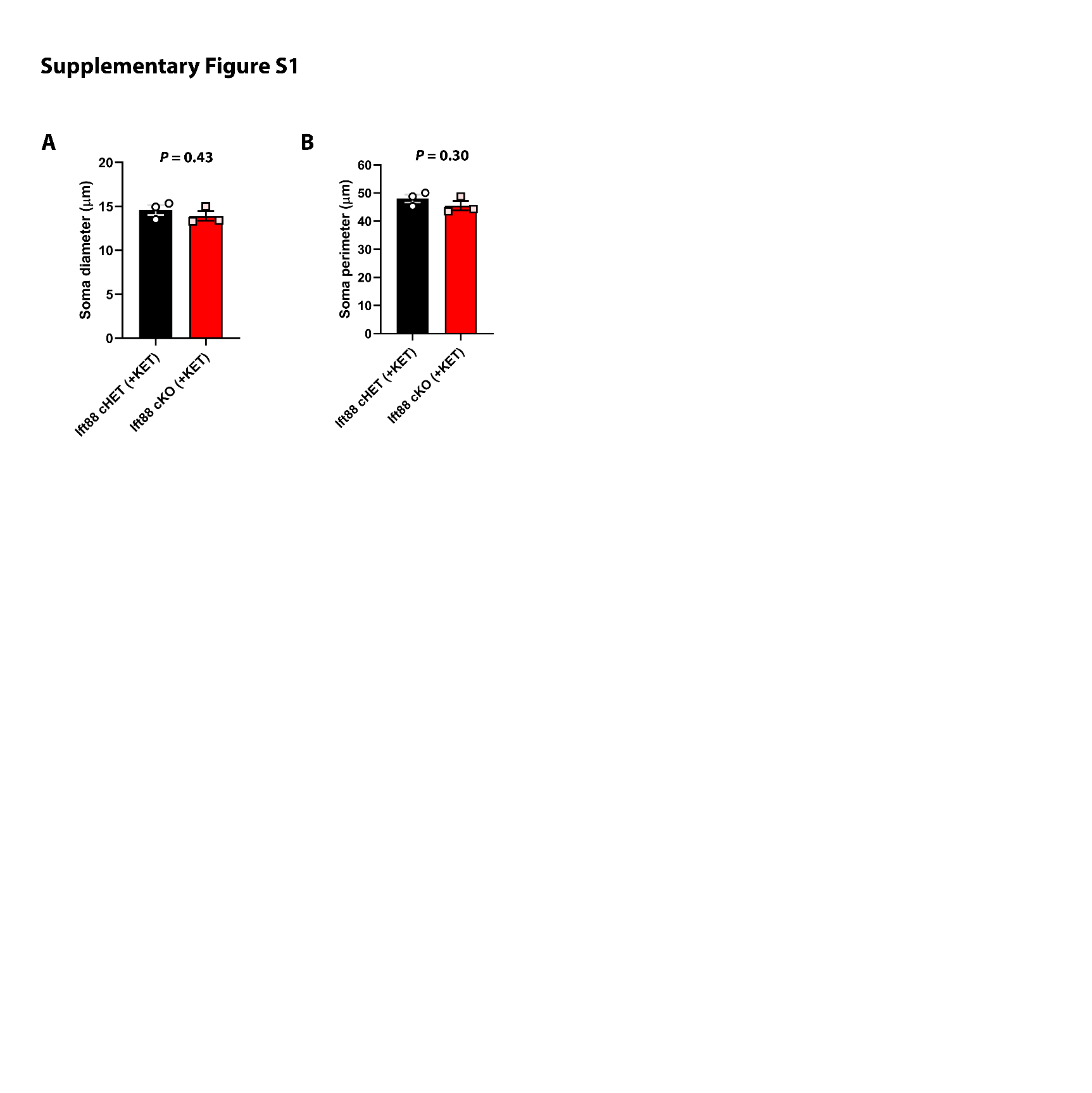

**Figure S1 Neuronal soma size does not change in ciliopathic motor cortical neurons exposed to ketamine.** Comparison of neuronal soma diameter (**A**) and perimeter (**B**) between adult (8-10 weeks) ketamine treated *Ift88* cHET and cKO mice. The statistical test used was a student’s unpaired t test.

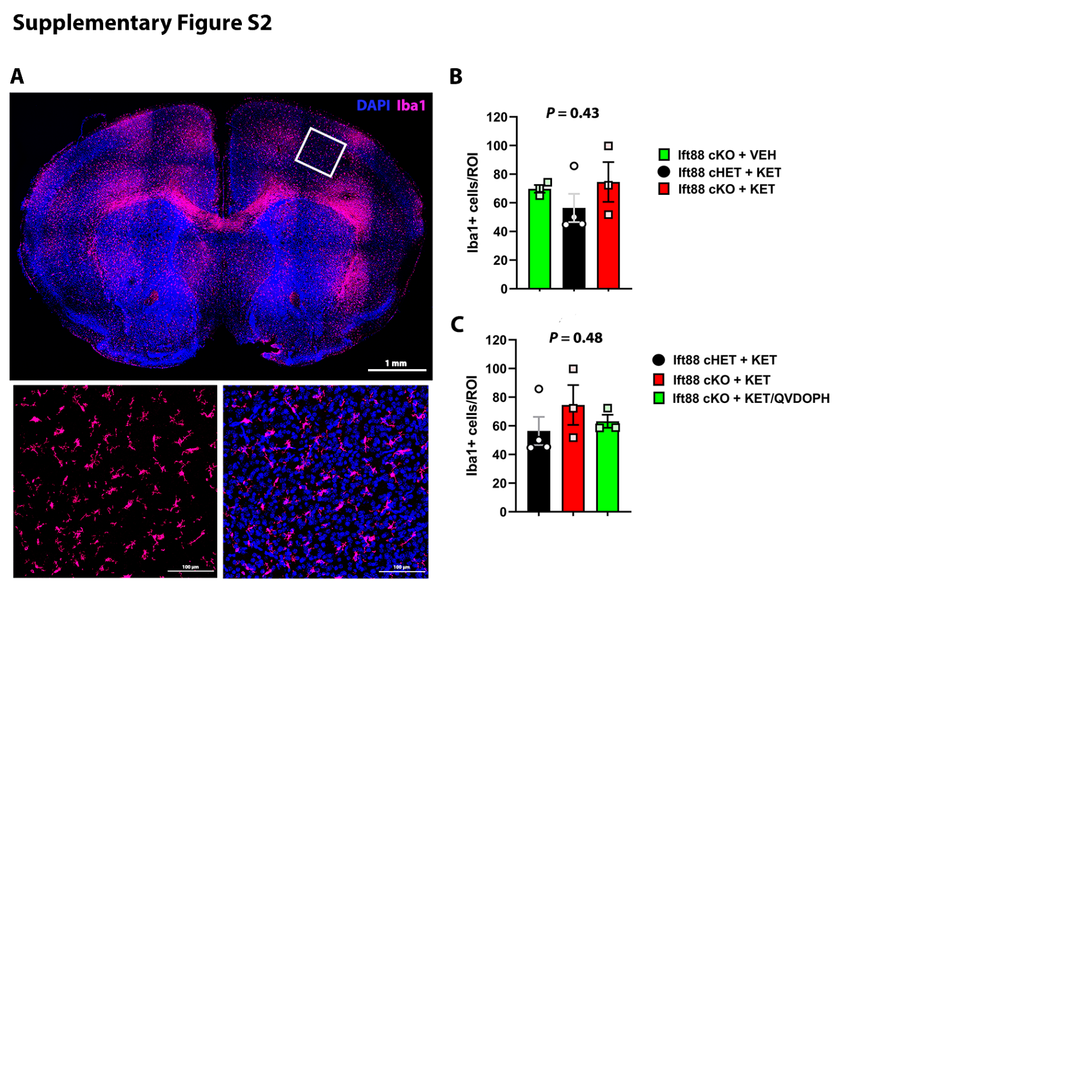

**Figure S2 Ketamine treatment does not elicit increased microglial infiltration in the ciliopathic motor cortex.** (**A**) Photomicrograph showing representative immunohistochemical staining for the microglial marker Iba1 at 16 hours post-ketamine treatment. (**B**) Quantification of Iba1^+^ microglia in the deep layers of the primary motor cortex (PMC). Note no significant difference in microglial numbers due to ketamine exposure. (**C**) Quantification of Iba1^+^ microglia in PMC deep layers indicates no differences following QVD-Oph treatment. Statistical tests used were one way ANOVA.

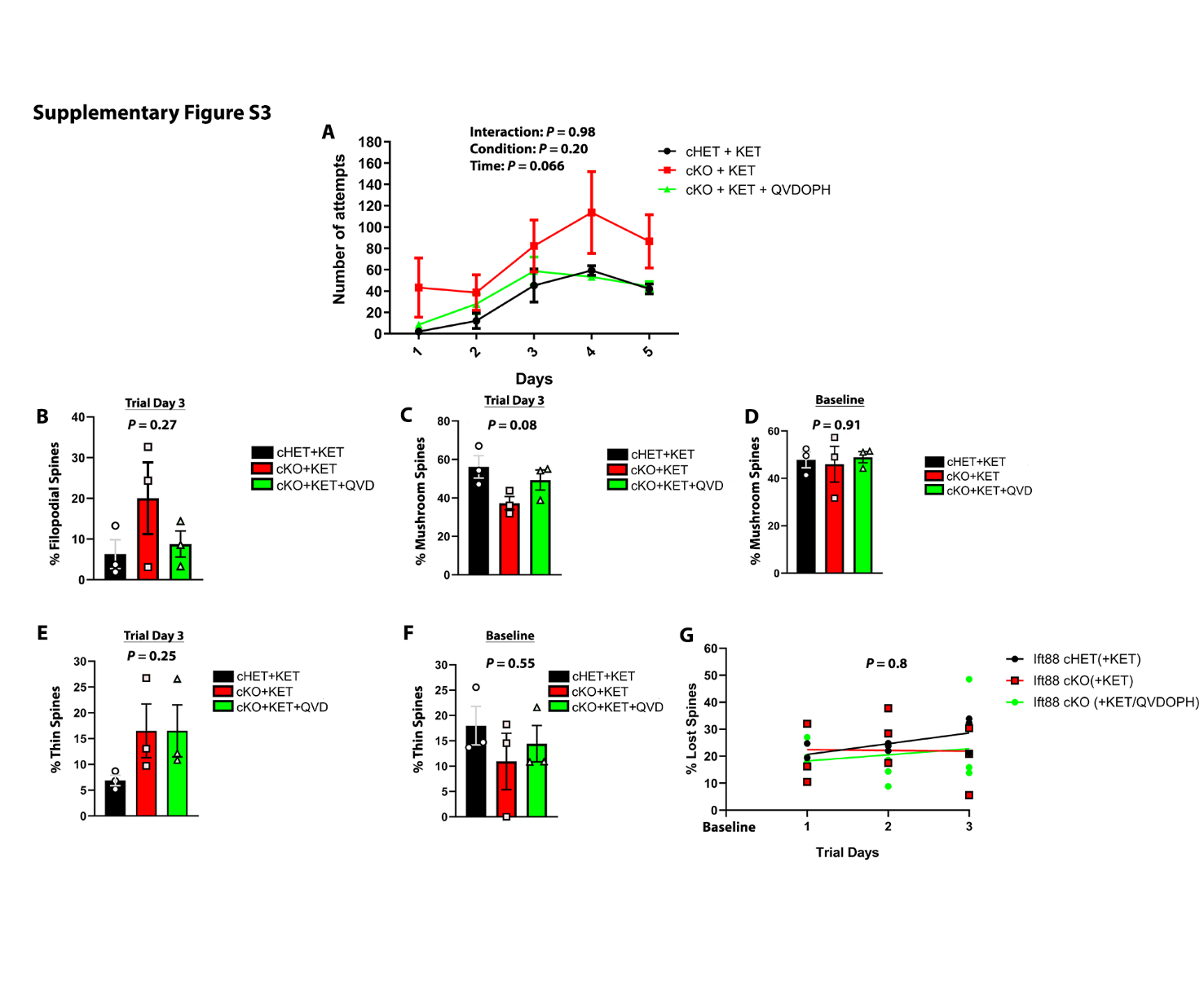
**Figure S3 QVD-Oph treatment effects on reaching behavior and dendritic spine class.** (**A**) Line graph showing total number of pellet reach attempts for groups assayed in Fig. 6B. Mice from all groups did not differ significantly in pellet reach attempts. Numbers used: ketamine treated *Ift88* cHET, *n* = 6; ketamine treated *Ift88* cKO, *n* = 9; ketamine + QVD-Oph treated *Ift88* cKO, *n*=5. The statistical test used was a 2-way repeated measures ANOVA with post-hoc Tukey’s multiple comparisons test. (**B**) Quantification of the fraction of filopodial spines at final training day. (**C-D**) Bar charts display fraction of mushroom/mature dendritic spines at final training day and baseline, respectively. Note statistical trend towards a rescue effect by QVD-Oph in the *Ift88* cKO group at the end of training. (**E-F**) Charts showing fractions of thin/immature spines detected in all groups at final training day and baseline, respectively. Note increase in thin spines on final training day in both *Ift88* cKO groups. Statistical tests used in (**C-F**) were one way ANOVA tests. (**G**) Line graph depicting the rate of dendritic spine loss during rotarod motor training in ketamine treated control (*Ift88* cHET) and forebrain-specific ciliopathy (*Ift88* cKO) groups given ketamine or ketamine with QVD-Oph. Note no significant difference in the rate of spine loss during motor training as assessed by linear regression. Values in all panels represent mean ± standard error (SEM).
